# Assessment of protein-protein interfaces in cryo-EM derived assemblies

**DOI:** 10.1101/2020.11.17.387068

**Authors:** Sony Malhotra, Agnel Praveen Joseph, Jeyan Thiyagalingam, Maya Topf

**Affiliations:** Institute of Structural and Molecular Biology, Department of Biological Sciences, Birkbeck College, University of London, Malet Street, London WC1E 7HX, United Kingdom; Scientific Computing Department, Science and Technology Facilities Council, Research Complex at Harwell, Didcot OX11 0FA, United Kingdom; Department of Engineering Sciences, Parks Road, Oxford, OX1 3PJ, United Kingdom; Centre for Structural Systems Biology, Heinrich-Pette-Institut, Leibniz-Institut fu□r Experimentelle Virologie and Universitätsklinikum Hamburg-Eppendorf (UKE), 22607 Hamburg, Germany

**Keywords:** Protein-protein complexes, cryo-EM, cryo-EM fitting, Assessment, Validation, Interface quality, Protein-protein interface

## Abstract

Structures of macromolecular assemblies derived from cryo-EM maps often contain errors that become more abundant with decreasing resolution. Despite efforts in the cryo-EM community to develop metrics for the map and atomistic model validation, thus far, no specific scoring metrics have been applied systematically to assess the interface between the assembly subunits. Here, we have assessed protein-protein interfaces in macromolecular assemblies derived by cryo-EM. To this end, we developed PI-score, a density-independent machine learning-based metric, trained using protein-protein interfaces’ features in high-resolution crystal structures. Using PI-score, we were able to identify errors at interfaces in the PDB-deposited cryo-EM structures (including SARS-CoV-2 complexes) and in the models submitted for cryo-EM targets in CASP13 and the EM model challenge. Some of the identified errors, especially at medium-to-low resolution structures, were not captured by density-based assessment scores. Our method can therefore provide a powerful complementary assessment tool for the increasing number of complexes solved by cryo-EM.

## Introduction

Proteins are known to interact with other biomolecules, and themselves, to perform their functions, and sustain the activities of cells. Unveiling the molecular details underlying these functions provides crucial structural, and functional insights. In recent years, cryo-EM has become a prominent technique for solving the structures of complex biological systems, such as polymerases, transmembrane receptors, viral assemblies and ribosomes, by overcoming some of the limitations of X-ray crystallography and NMR spectroscopy^1^. Cryo-EM techniques, which usually require a small amount of sample, are more forbearing on sample purity, and the rapid freezing of the sample maintains its closeness to native state. Due to these strengths, cryo-EM provides an alternative to X-ray crystallography for large complexes. Recent advances in instrumentation and image processing methods in structure determination using cryo-EM and tomography of sub-cellular structures have pushed the resolution of structures. However, the average resolution of structures solved using single-particle cryo-EM every year is worse than 5 Å (5.6 Å for 2019 and 6.2 Å for 2020), and determining structures at near-atomic resolution is still a challenge and largely sample dependent^2,3^. Additionally, many of the cryo-EM maps associated with a near-atomic global resolution have regions at intermediate resolutions (or even lower), owing to the variability in local resolution.

The resolution of the cryo-EM map dictates the approach to be adopted for model building, fitting, refinement and validation to a great extent^4^. Regardless of the resolution of the map, upon model building and/or fitting, assessment of the atomistic model is crucial to ensure its overall reliability, and thus should be independent of the score(s) optimised during the fitting stage.

The most commonly used global score to optimise the fitting is density cross-correlation coefficient (CCC) between the cryo-EM map, and simulated density of the fitted atomistic model. Apart from some variations of the CCC with masks and filters, there are other global scores, such as the mutual information^5^. Local scores are very useful in identifying the regions of poor fit in the models, which can be further refined to obtain a better fit. Local Mutual Information (MI), TEMPy local scores-SMOC^6^ (segment-based Manders’ overlap coefficient) and SCCC^7^ (segment-based cross correlation score), Q-scores^8^, and EMRinger^9^ are scores that can guide the fitting at different structure representations, such as residues, domains, secondary structure elements, and loop regions. Additionally, there are other metrics that assess the geometry of models, such as MolProbity^10^ and CaBLAM^11^. These metrics, however, do not include the assessment of the quaternary structure in terms of quality of the interface between subunits.

Some of the common scenarios that may result in sub-optimal protein-protein interfaces in cryo-EM derived models are as follows:

⍰ fitted models are usually built sequentially, *i*.*e*. one chain is fitted into the map at a time, independent of the others;
⍰ map segmentations are an integral part of model building, but segmentation techniques are not accurate enough to identify boundaries between the subunits;
⍰ building the model of only one protomer and applying symmetry operations; and
⍰ integrating models of subunits built in maps reconstructed by refinement focused on certain segment(s) of the macromolecule.

Scoring quality of protein-protein interfaces will provide crucial model quality assessment, especially for the cases listed above. The features that characterise the interfaces can be used to build an overall metric, and while being density-independent, to provide an additional complementary quality measure to assess the quality of the modelled assembly in the cryo-EM map. Features that have been shown to be discriminatory in identifying biological (*‘native-like’*) interfaces are, for example, conservation of residues present at interface^12–16^, shape^17,18^ and electrostatic complementarity^19,20^, residue contact pairs^21,22^, types of interactions ^23–26^, and interface size and area^25,27–29^.

Although these features are useful in identifying the quality of interfaces, these derivations are often done in isolation, and largely relies on the expertise of the scientists for validation based on their experience. As such, computational algorithms for computing scores based on the metrics and features described above, ignores one major aspect – extraction and reuse of the knowledge from different datasets. Machine learning (ML)-based approaches, on the other hand, are inherently data-centric, can accumulate knowledge from various datasets. ML, a class of algorithms that learns from the data are often *trained* on several datasets prior to using them on real datasets (inferencing). At the simplest level, the model learns to classify and distinguish *‘native’* and *‘native-like’* interactions from the false interactions.

In fact, a number of approaches have utilised ML methods for predicting protein-protein interactions. These approaches vary in terms of exact algorithms used, datasets (*i*.*e*., protein-protein complexes), and more importantly, on the set of features used for training. The most commonly used features for predicting protein-protein interactions using ML-based methods include physicochemical properties, evolutionary features, secondary structures, solvent accessible area and binding energies among others. The choice of algorithms for training ML models include Support Vector Machines (SVM), Random Forest (RF), Neural Networks (NN) or ensemble learning^30^. Combination of different features and ML algorithms lead to a very rich set of methods that one can rely on. Recent reviews^30,31^, provide an elaborative comparison of structure- and sequence-based existing methods, providing detailed account on their performance and availability of those techniques.

In this article, we present a systematic assessment of protein-protein interfaces in cryo-EM derived assemblies using a new metric-Protein Interface-score (PI-score). The score was developed based on various features describing protein-protein interfaces in high-resolution crystal structures from the Protein Data Bank (PDB). These derived features were further used to train a ML-based classifier in order to distinguish ‘good’ (native/native-like) and ‘bad’ interfaces. To assess the applicability and performance of the trained model to cryo-EM derived assemblies, we used PI-score to assess the quality of interfaces in CASP-13 cryo-EM targets, EM model challenge targets (2016 and 2019), and PDB entries associated with Electron Microscopy Data Bank (EMDB) (4913, as of Aug 2020).

## Results

In this section, we discuss the workflow of building a training and testing dataset for a machine learning (ML)-based model to assess protein-protein interfaces. The derived score (PI-score) is then applied to assess the quality of interfaces in models submitted for CASP13 targets, EM model challenge targets, and PDB entries associated with EMDB. We discuss the examples from each of these datasets. We also compared the performance of PI-score with other density based-score and statistical interface potentials.

### Building the Dataset

A total of 3,926 high-resolution complexes obtained from PDB^32^ were subjected to an *in-house* pipeline to assign the interfaces using a distance-based threshold (Methods). To avoid the over-representation of similar interfaces in the dataset, structurally similar interfaces within a quaternary structure were filtered out using interface similarity score calculated with *iAlign*^33^, resulting in 2,858 interfaces from 2,314 complexes. Various interface features, namely: number of interface residues, contact pairs, surface area, shape complementarity, number of hydrophobic, charged, polar and conserved residues at the interface and other interface properties evaluated by PISA, were computed for the dataset. These features were successfully calculated for 2,406 interfaces, which form “positive dataset1” (PD1, see Methods).

To train the ML classifiers of our choice on data closer in quality to models fitted on cryo-EM maps (especially at intermediate-to-low resolutions), noise was added to PD1 by slightly perturbing the relative positions and orientations of the interacting subunits. This was performed using a protein-protein docking method^34^ and then selecting the poses with high fraction of aligned interface residues/interface residues in native complex (f_Nal_), and low interface RMSD (iRMSD) (see Methods for cut-offs). This set, which contains 3,743 interfaces, is referred to as “positive dataset2” (PD2).

A “negative dataset” (ND), containing 3,578 interfaces, was also derived using docking and includes complexes in which the interfaces are structurally different, *i*.*e*., ‘far’ from native interfaces (low f_Nal_ and high iRMSD, see Methods for cut-offs). The schematic of the procedure to collate the datasets and workflow is summarised in Figure 1.

**Figure 1:**
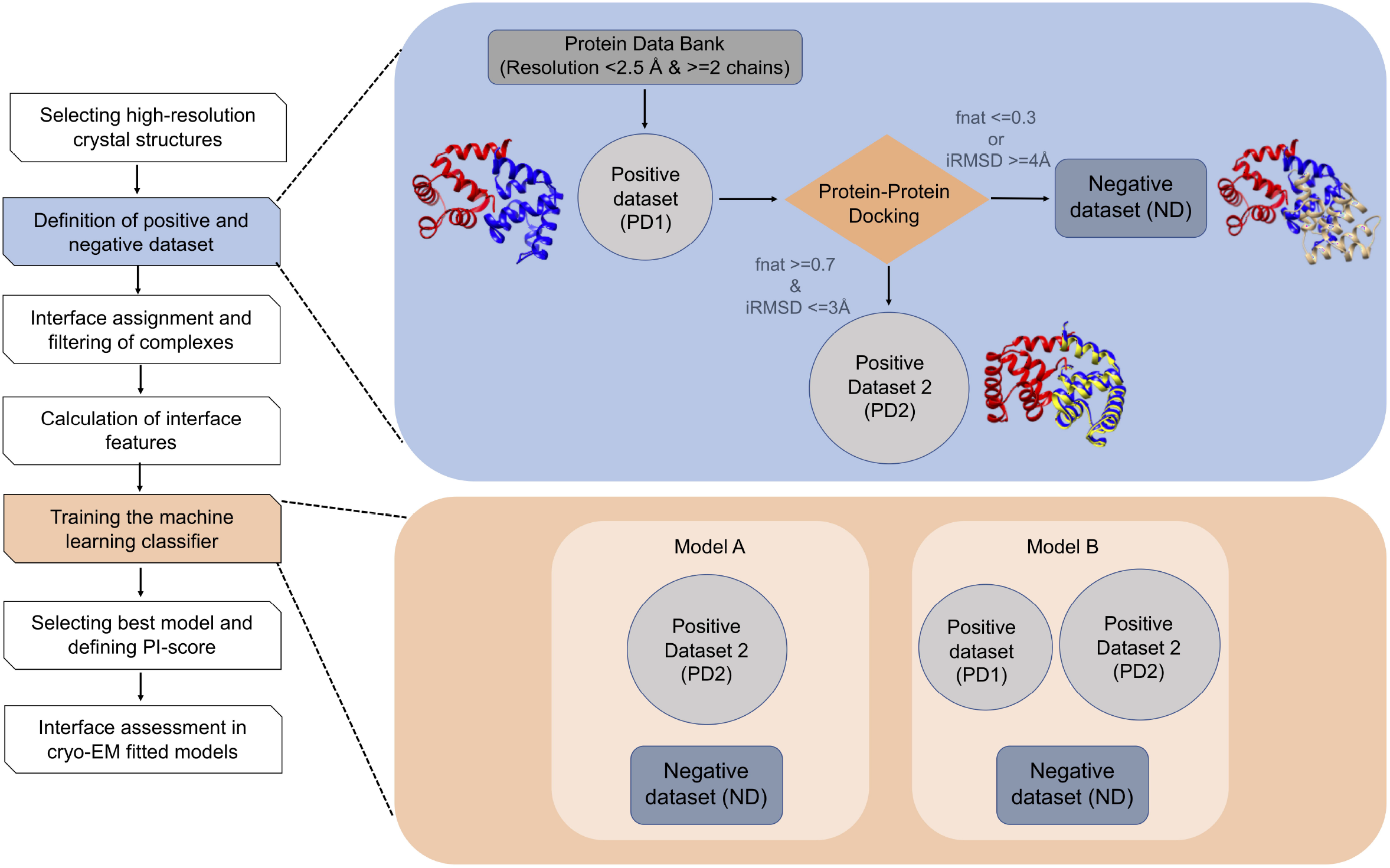
Workflow for developing a protein-protein interface-based score (PI-score) to assess macromolecular assemblies derived using cryo-EM. High resolution complexes (with >=two chains) were obtained from the PDB and are referred to as the “positive dataset1” (PD1). Protein-protein docking was used to derive, structurally close (to PD1) complexes to form the “positive dataset2” (PD2). The complexes obtained upon docking that have a higher interface RMSD (iRMSD) and lower fraction of aligned native residues (f_Nal_) at the interface, form the “negative dataset” (ND). Interface features were calculated on all the complexes and are used as an input to train a supervised machine learning classifier, which is further used to predict the class labels of the benchmark dataset.

### Ranking of Interface Features

Various interface features (listed in Methods) were computed for the above-described datasets. As the number of derived features (12) was manageable and computationally not very expensive, we used all the features to train our classifiers. To identify the top-ranking (or most influential) interface feature(s), we used different methods namely, Ridge, Random Forest, Recursive Feature Elimination, Linear Regression and Lasso for feature ranking. Our exploration showed that the top-ranking features were shape complementarity, number of polar interface residues, number of charged interface residues, and interface solvation energy (Figure 2a).

**Figure 2:**
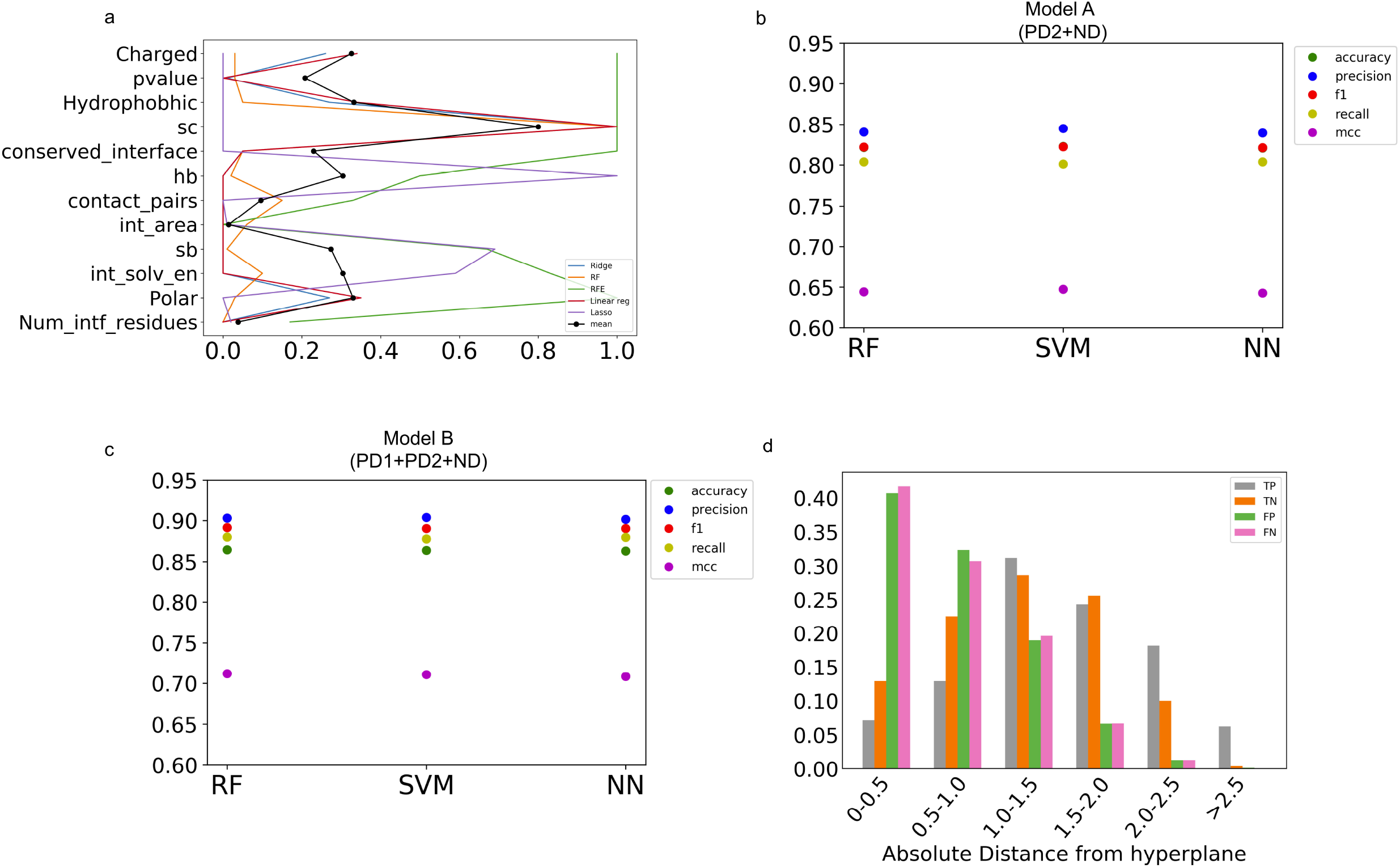
Machine learning based classifier to assess the quality of protein-protein interfaces. a) Importance of interface features in distinguishing the ‘native-like’ interface. The ranks from different methods (Ridge, Random Forest (RF), Recursive feature elimination (RFE), Linear regression (Linear reg) and Lasso) were normalised between 0-1 and the mean feature rank is plotted in black. b) and c) Performance of different classifiers on the training dataset: RF (Random forest), SVM (Support vector machine), and NN (Neural Networks) are used to perform supervised learning on the training dataset using *stratified shuffle split* as a means of cross-validation with ten splits. The performance is evaluated using accuracy, precision, F1, recall scores and Matthews correlation coefficient. Performance measures of Model A (b): trained on docking-derived positive dataset (PD2) and negative dataset (ND). Performance measures of Model B (c): trained using both high-resolution and docking-derived positive datasets (PD1+PD2) and negative dataset (ND). d) Fraction of True Positives (TP), True Negatives (TN), False Positives (FP) and False Negatives (FN) in different PI-score thresholds. The fractions (Y-axis) are averaged over the ten splits (*stratified shuffle split*) of the data. The different PI-score thresholds (X-axis) are indicated in absolute values.

### Training the Classifier and Cross-Validation

To develop a better understanding of the performance, we evaluated three ML classifiers, namely, Support Vector Machine (SVM), Random Forest (RF), and plain vanilla Neural Network (NN), or simply, multi-layer perceptron (MLP), using the Scikit-learn Python package^35^ (*scikit*).

We used the following combination of datasets described above to train two high-level models, namely, Model A and Model B (referred to as models henceforth), using the interface features (Methods) (Figure 1). Each of these models relies on three different classifiers, described above (SVM, RF and NN).

### Model A

Positive and negative datasets derived using docking (PD2 and ND, respectively).

### Model B

Positive dataset constitutes high-resolution complexes and computationally derived docked complexes (PD1 + PD2) and ND as negative dataset.

While training and testing both models (Model A and Model B), to minimise the bias of the classifiers, which can easily become an issue with unbalanced datasets, we used *stratified shuffle split* with ten splits (and test size of 30%) as a measure of cross-validation. In both scenarios, the performance, which was measured based on ML- and classifier-specific metrics, namely, accuracy, precision, recall, F1 and Matthew’s Correlation Coefficient (MCC), was comparable between the three methods (Figure 2b and Figure 2c).

Among these, the SVM-trained model offered the best validation accuracy of 86% (Model B), and it was selected to assess the quality of protein-protein interfaces modelled in cryo-EM maps. The SVM classifier finds a hyperplane that maximises the inter-class variance, and we used the distance of a given data-point from this hyperplane as the machine learning-based score (PI-score) for a given prediction (interface). The farther a point (interface) is located from the hyperplane (more negative or positive), the more confident is the prediction using the SVM model^36^. We assessed the performance of the PI-score at different thresholds by analysing the number of false positives (FP) and false negatives (FN). For the ten test sets (30%) obtained using the stratified shuffle split (for cross validation purpose), the fractions of FP (41%) and FN (42%) were observed to be highest in the PI-score ranges of (0 to 0.5] and (−0.5 to 0], respectively (Figure 2d). We also estimated the measures of performance in different PI-score thresholds and observed that the PI-scores >1 and <-1 (for the positive and negative class label, respectively), were more reliable, based on the low false positive rate (FPR) and high true positive rate (TPR) in the respective bins (Table 1).

**Table 1:** Performance in different bins of the scores using the SVM machine learning-based classifier. The following bins according to the listed thresholds and the measure TPR, FPR, Precision and Specificity are averaged values over the test datasets obtained from ten-fold cross-validation: 0.0-0.5: True positives present in the score range of 0.0 to 0.5 and true negative in score range of −0.5 to 0.0. False positives are the complexes from negative dataset (ND) but are predicted positive with a score assigned between (0.0 to 0.5) and false negatives are positive complexes from either positive dataset 1 or 2 (PD1 or PD2) but are predicted as negative with the score between (−0.5 to 0.0). 0.5-1.0: True positives present in the score range of 0.5 to 1.0 and true negative in score range of −0.5 to −1.0. False positives are the complexes from negative dataset (ND) but are predicted positive with a score assigned between (0.5 to 1) and false negatives are positive complexes from either positive dataset 1 or 2 (PD1 or PD2) but are predicted as negative with the score between (−0.5 to −1.0). 1.0-1.5: True positives present in the score range of 1.0 to 1.5 and true negative in score range of −1.0 to −1.5. False positives are the complexes from negative dataset (ND) but are predicted positive with a score assigned between (1.0 to 1.5) and false negatives are positive complexes from either positive dataset 1 or 2 (PD1 or PD2) but are predicted as negative with the score between (−1.0 to −1.5). 1.5-2.0: True positives present in the score range of 1.5 to 2.0 and true negative in score range of −1.5 to −2.0. False positives are the complexes from negative dataset (ND) but are predicted positive with a score assigned between (1.5 to 2.0) and false negatives are positive complexes from either positive dataset 1 or 2 (PD1 or PD2) but are predicted as negative with the score between (−1.5 to −2.0). 2.0-2.5: True positives present in the score range of 2.0 to 2.5 and true negative in score range of −2.0 to −2.5. False positives are the complexes from negative dataset (ND) but are predicted positive with a score assigned between (2.0 to 2.5) and false negatives are positive complexes from either positive dataset 1 or 2 (PD1 or PD2) but are predicted as negative with the score between (−2.0 to −2.5). >=2.5 and <=-2.5: True positives with a score >= 2.5 and true negative with score <= −2.5. False positives are the complexes from negative dataset (ND) but are predicted positive with a score >2.5 and false negatives are positive complexes from either positive dataset 1 or 2 (PD1 or PD2) but are predicted as negative with the score < −2.5.

| Scores' bins | TPR (True Positive Rate) | FPR (False positive Rate) | Precision | Specificity |
| --- | --- | --- | --- | --- |
| (-/+)[0.0 to 0.5) | 0.15 | 0.76 | 0.15 | 0.24 |
| (-/+)[0.5 to 1.0) | 0.29 | 0.58 | 0.29 | 0.41 |
| (-/+)[1.0 to 1.5) | 0.61 | 0.39 | 0.63 | 0.61 |
| (-/+)[1.5 to 2.0) | 0.78 | 0.21 | 0.78 | 0.78 |
| (-/+)[2.0 to 2.5) | 0.93 | 0.11 | 0.93 | 0.99 |
| $\geq 2.5$ and $\leq -2.5$ | 1.0 | 0.18 | 0.97 | 0.81 |

### Application to CASP Targets (High Resolution Targets)

We applied the above trained models to make predictions on the quality of protein-protein interfaces in cryo-EM targets from the CASP13 competition^37^. Three of the targets (T1020o, T0995o, and T0984o) were classified as ‘easy targets’, with many high-accuracy models deposited by the participating groups and were also evaluated for the goodness-of-fit to the experimental cryo-EM maps^38^. For each of these targets, a submitted pool of models, an experimentally solved structure (target) and a density-based score for assessing the goodness-of-fit are available from the CASP13 website. Therefore, these targets form an ideal dataset for assessing the performance of PI-score.

We used the CASP multimeric scores (https://predictioncenter.org/casp13/multimer_results.cgi), namely, F1, Jaccard index, lDDT(oligo) and GDT(o) (see Methods: comparison with CASP13 oligomeric scores) to define true positives (TP), true negatives (TN), FP and FN for CASP targets. If any of the four CASP13 multimeric scores was equal or greater than (>=) 0.5, and the model was scored positive by our classifier, it was treated as TP. TN were defined as model structures which did not have any of the CASP13 scores >=0.5 and were scored negative by the classifier. The models which were scored >=0.5 by any of the four CASP13 scores and negative using our classifier score were defined as FP and models which were scored negative by the classifier but had at least one of the CASP13 score >=0.5 were FN.

### T1020o: 3.3 Å resolution homo-trimer structure of an S-type anion channel from Brachypodium distachyon

Using our SVM classifier, nine of the assessed 111 submitted models (with 329 interfaces) were predicted to have at least one ‘negative’ interface (negative PI-score) in the complex. These nine models were also scored low on the CASP multimeric assessment scores^39^ (Supplementary Table S1). With a more systematic comparison of PI-scores against the oligomeric assembly assessment scores from CASP13^39^, we achieve 82% accuracy for this target (see Methods: comparison with CASP13 oligomeric scores).

All interfaces in the target structure (Figure 3a) and in the top-ranked model based on the cross-correlation of the model with the cryo-EM density (CCC) ((TS004_2o, Figure 3b) have positive PI-score. Out of the nine negatively-scoring models, TS008_4o and TS135_3o had negative PI-score for all the three interfaces (Figure 3c and Figure 3d, respectively). When these models are compared to the target structure, all three interfaces have high iRMSD and low fraction of aligned native residues (Average iRMSD of 2.93 Å and 3.33 Å for TS008_4o and TS135_3o, respectively, Table 2). For model TS208_1o, two of the interfaces (formed by chains, AC and BC) have negative PI-score (Figure 3e) and PI-score was not calculated for the third interface, as the number of interface residues were only nine and eight for chain A and B, respectively, which was less than our cut-off for defining an interface (Methods).

**Table 2:** Assessment of interfaces in the models of CASP13 cryo-EM target T1020o. The model and equivalent target chains forming the interface are listed along with the interface RMSD (iRMSD), fraction of aligned native interface residues (f_Nat_) and predicted class using our model.

| Model ID | Model interface | Target interface | iRMSD (Å), $f_{\text{Nat}}$ | Predicted class | Score |
| --- | --- | --- | --- | --- | --- |
| TS004_2o | AB | AB | 2.2, 0.81 | Positive | 2.6 |
|  | BC | BC | 2.5, 0.75 | Positive | 2.6 |
|  | AC | AC | 2.1, 0.82 | Positive | 2.7 |
| TS008_4o | AB | AC | 3.16, 0.42 | Negative | -1.5 |
|  | BC | BC | 2.82, 0.48 | Negative | -1.5 |
|  | AC | AB | 2.81, 0.48 | Negative | -1.5 |
| TS135_3o | AB | BC | 3.08, 0.56 | Negative | -1.6 |
|  | BC | AB | 3.4, 0.52 | Negative | -1.6 |
|  | AC | AC | 3.51, 0.6 | Negative | -1.6 |
| TS208_1o | AB | BC | 2.6, 0.63 | Not ranked<br>(Interface residues from model 9 and 8) | NA |
|  | BC | AC | 2.5, 0.54 | Negative | -0.2 |
|  | AC | AB | 2.6, 0.52 | Negative | -0.39 |

**Figure 3:**
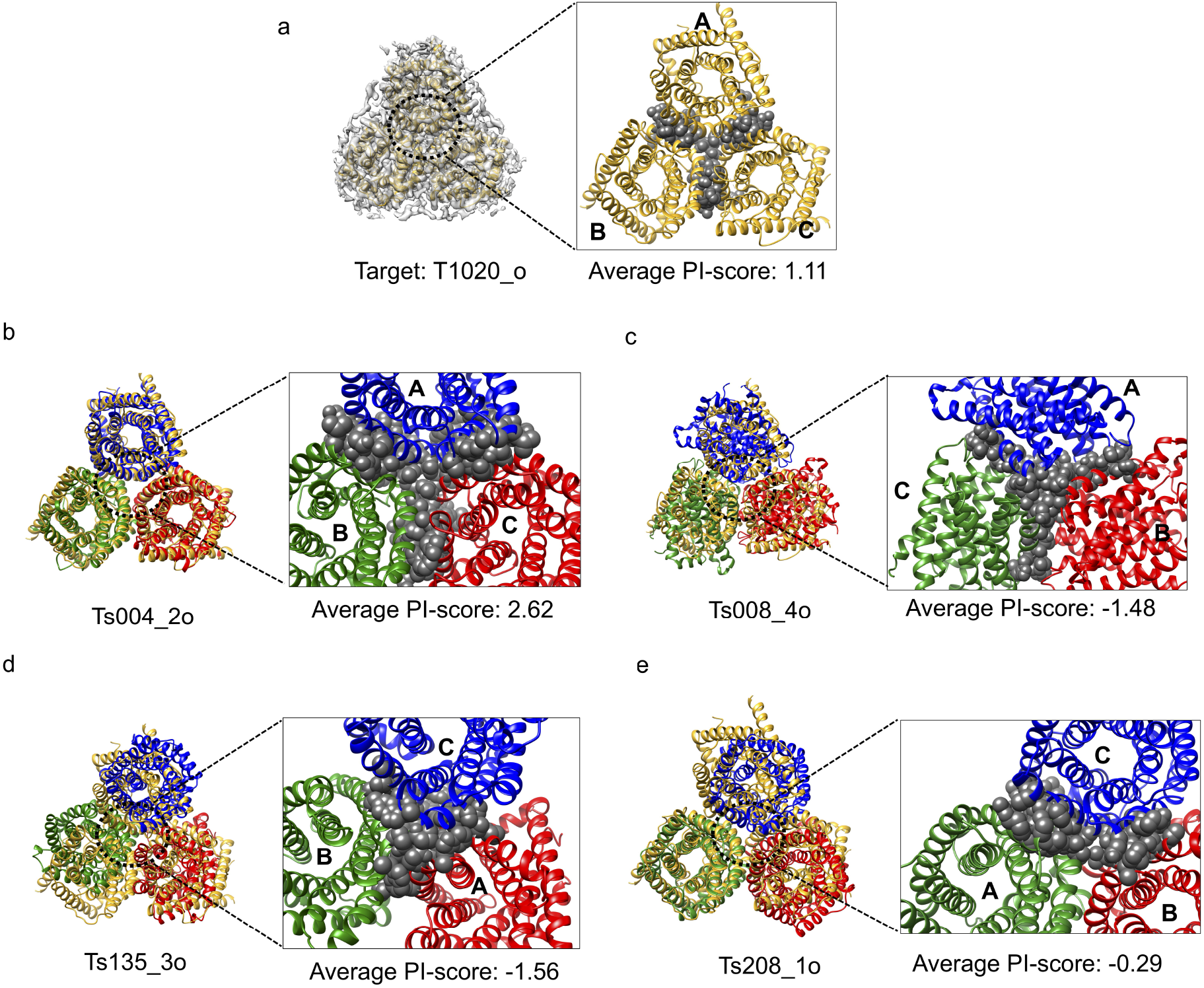
Scoring the interfaces in the oligomeric CASP13 target T1020o. The target structure is shown in gold in all the panels and the model structures being assessed are shown in red, green and blue. The chains are labelled accordingly. a) Target structure within the cryo-EM map. The interface residues from the three chains are shown as grey spheres. b) Model TS004_2o, with a positive PI-score for all the three interfaces in the trimeric assembly. c), d) and e) Models TS008_4o, TS135_3o and TS208_1o, respectively, for which their interfaces are scored negatively with PI-score.

For the model TS208_1o (CCC = 0.42, target structure CCC = 0.77), we generated density maps at resolutions lower than the target map: 5, 8, 10 and 12 Å using the ‘*low pass filter’* utility in CCP-EM suite (https://www.ccpem.ac.uk/). Since the CCC does not have a defined absolute cut off value to differentiate between good and bad fit at any given resolution, it is difficult to identify ‘target-like’ models. On the other hand, PI-score, which is a density-independent metric, can be very useful to distinguish ‘target/native-like’ interfaces in the modelled complex(es) (Supplementary Table S2).

### T0995o: A 3.15 Å resolution homo-octamer (A8) of cyanide dehydratase

The PI-score for the target structure was positive for the dimer interface (Supplementary Figure S1a), which is repeated to form an octameric complex. We calculated the PI-score for 657 interfaces in the 118 CASP13 models for this target and assessed the quality of the dimer interface between all subunits. 123 interfaces in 37 models were observed to have negative PI-score. The top-ranked model (after target) in terms of CCC was TS008_2o (Supplementary Figure S1b), which is calculated to have positive PI-score for the equivalent dimer interface (iRMSD=1.55 Å). Examples for the models with negative PI-score are TS117_1o (iRMSD = 4 Å, Supplementary Figure S1c) and TS008_5o (iRMSD = 2.76 Å, Supplementary Figure S1d). The models with interfaces having negative PI-score using our classifier were also scored low for the CASP13 multimeric scores (Supplementary Table S1).

By comparing it with the multimeric scores in CASP13, we achieve an accuracy of 67% for this target. This target has higher stoichiometry and more interfaces than T1020o, and therefore it is expected to achieve a lower accuracy against the CASP13 assembly scores, which are calculated per complex (while our classifier is per interface and hence this may not be a direct comparison).

PI-scores for the assessed interfaces in models for CASP13 cryo-EM targets is provided in Supplementary Table S3.

### T0984o: A 3.4 Å dimer of a calcium channel

145 models were assessed, and all were observed to have a positive PI-score for the interface (Supplementary Table S3).

Given the nature of CASP experiments where the participating groups model the complexes without the knowledge of cryo-EM map, protein-protein interface assessments such as PI-score will provide complementary assessment and are crucial to provide insights into model quality.

### Application to EM Model Challenge

Next, we calculated PI-scores for the models submitted for the targets from two EM validation challenges (https://challenges.emdataresource.org/), namely, 2016 EM model challenge and 2019 model metrics challenge (Supplementary Table S4).

Target T0002 (from model challenge 2016) is a 3.3 Å resolution cryo-EM map of the 20S proteasome (EMD-5623). We assessed the ten submitted models (with 175 interfaces) and interfaces in the target structure (PDB ID: 3J9I). In three of the models-EM164_1, EM189_1 and EM189_2, there was at least one interface that obtained a negative PI-score.

As an example, we chose model EM164_1, for which most the interfaces in the alpha and beta subunits were scored negative (Supplementary Table S4). In the alpha ring, the two subunits in the model (chains F and C, shown in red and green Figure 4a) were scored negative by our classifier (PI-score: −2.27). The interface conformation is slightly different as compared to the target structure (iRMSD = 0.86, f_Nal_ = 0.54). This interface is loosely packed (Figure 4) and smaller than the equivalent interface in the target structure (23 interface residues in model and 37 in the target structure). Due to its small size the iRMSD is low, and therefore is not a good indicator of the quality of the modelled interface in this case. Due to the offset in the modelled interface the shape complementarity at the interface drops significantly to 0.32 as opposed to 0.73 for the interface in the target structure. We also checked the multimeric scores from CASP assessment and EM164_1 is scored high for QS-global (0.88) and lDDT (0.98)) scores but these scores reflect the quality of a multimeric structure as a whole rather than per interface. Other interface assessment scores (from CASP13) such as F1 and Jaccard index, which are calculated per interface, are not reported in the CASP website for this model.

**Figure 4:**
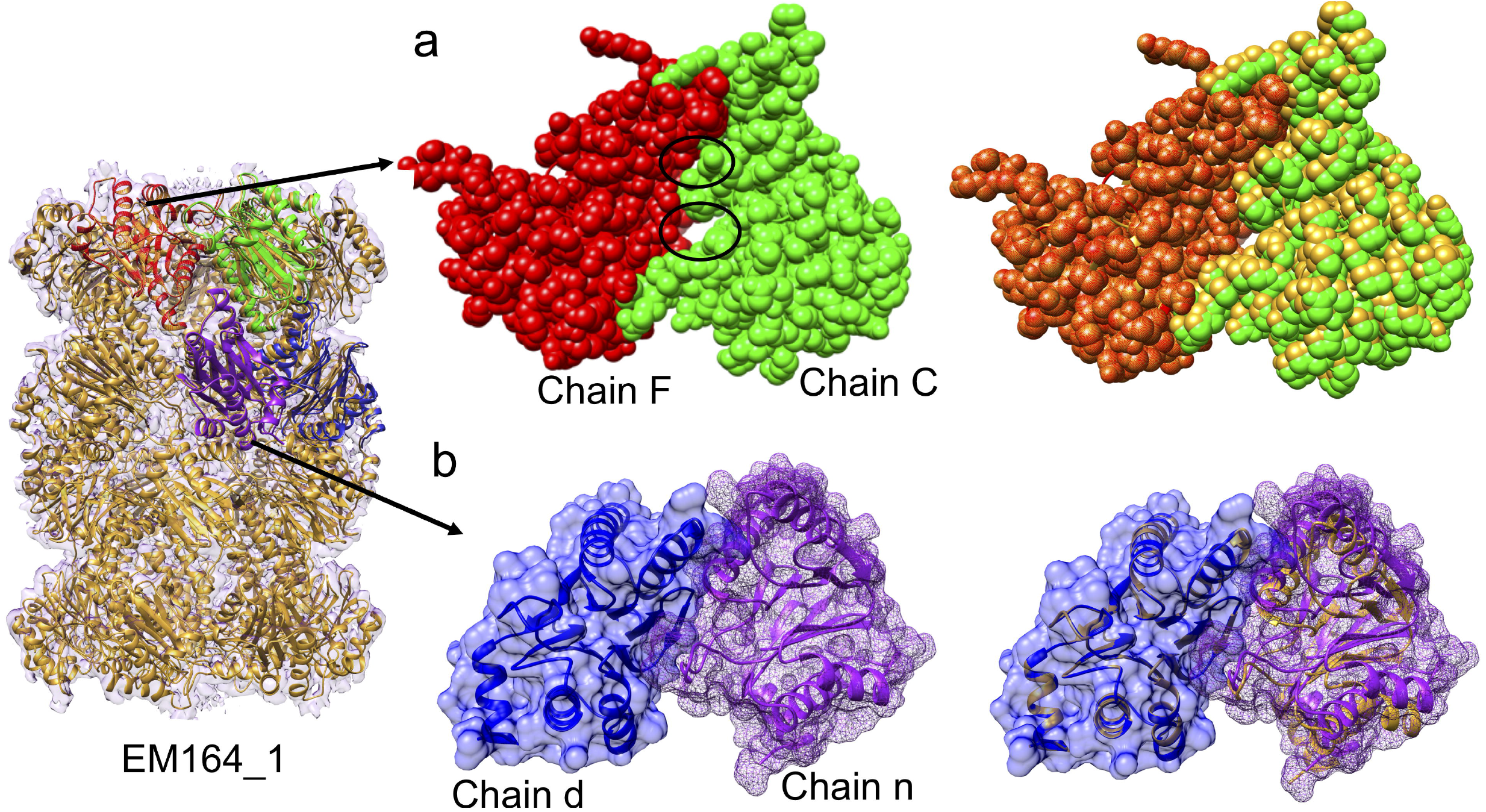
Scoring the interfaces in the target T0002 from 2016 EM model challenge. a) Scoring the interfaces between the alpha subunits’ ring in the model structure EM164_1, the chains (F and C), which form a negatively-scoring interface is shown in red and green and the target structure is shown in golden. The surface for the interface forming chains (F and C) are shown as spheres and the loose packing at the interface is marked with black ovals. b) Scoring the interfaces in the beta subunits’ ring of the 20S proteasome, one of the targets in EM model challenge (T0002). The target structure is shown in golden and the chains forming the interface (n and d) in the model structure (EM164_1) being assessed are shown in blue and purple. The surface (mesh) of the chains forming interface are shown to highlight the clashes at the interface formed by chains n and d in the model.

Structurally equivalent subunits in the target structure (chains P and Q) have a CCC of 0.85 and the model subunits (chains F and C) have a CCC of 0.73 (Supplementary Table S5) and local score (SMOC) averaged over interface residues are 0.73 and 0.23 for target and EM164_1 respectively. Our model has rightly predicted this interface as ‘negative’ as reflected by the loose packing at the interface and lower local density-based score for the modelled interface.

Further, we calculated the density-based scores (global and SMOC) at different resolutions (map simulated using *low pass filter utility* in CCP-EM, Supplementary Table S5). The scores assessing the fit of the model (with interface offset) are comparable at resolution worse than 5 Å. Therefore, especially at intermediate-low resolution, our proposed density-independent PI-score can be a crucial model validation tool.

The interfaces between the beta-subunits were also scored negative (TS164_1) by our classifier. This is reflected by the presence of steric clashes at the interface (blue and purple in Figure 4). The clashes present at the interface resulted in lower shape complementarity score for the interface in model (0.28) as opposed to a higher score (0.62) for the equivalent interface in the target structure. The subunits (chains X and Y) have a CCC of 0.85 whereas the subunits (chains n and d) of model have a CCC of 0.65. This model interface also has a much lower SMOC score than the equivalent interface in the target at all resolutions (Supplementary Table S5).

Recently, model metrics challenge (2019) was open, and we applied our score for assessing the only multimeric target −T0104 (Horse Liver Alcohol Dehydrogenase, 2.9 Å, dimer). We assessed the reference structure (PDB ID: 6NBB) and 17 submitted models using PI-score. Two models (T0104EM060_1; PI-score −0.31 and T0104EM060_2; PI-score 0.13), were scored low (Supplementary Table S4), which is in agreement with the CASP multimeric scores (QS and lDDT scores, https://challenges.emdataresource.org/?q=model-metrics-challenge-2019).

### Application to Fitted Entries in EMDB

We divided this dataset into three sets: high resolution (better than 4Å, ‘high resolution’), 4-8 Å (‘intermediate resolution) and 8-12 Å (low resolution). As we have described above the performance of PI-score using high-resolution complexes from CASP and the EM model challenge targets, in this section we will focus more on selected examples from intermediate and low resolution cryo-EM maps. The fitted models were also compared with the interfaces in the corresponding crystal structures.

For completeness, we also provide the PI-scores of our SVM model for the interfaces fitted at high resolution (better than 4Å) in Supplementary Table S6.

### Intermediate resolution (4-8Å)

Chikungunya virus: A cryo-EM map resolved at 5 Å (EMD-5577) with fitted model is available for the virus (PDB: 3J2W, shown in green and red in Figure 5a, with interface residues shown as grey circles). The envelope1-envelope2 (E1-E2) heterodimer was observed to have a negative PI-score (−1.67). The available crystal structure (PDB: 3N44, 2.35Å) for the E1-E2 subcomplex (chains B and F, coloured in golden and interface residues in grey spheres, Figure 5a) is scored positive (PI-score: 1.67). The interface between E1-E2 is slightly shifted as compared to the crystal structure (Supplementary Table S7).

**Figure 5:**
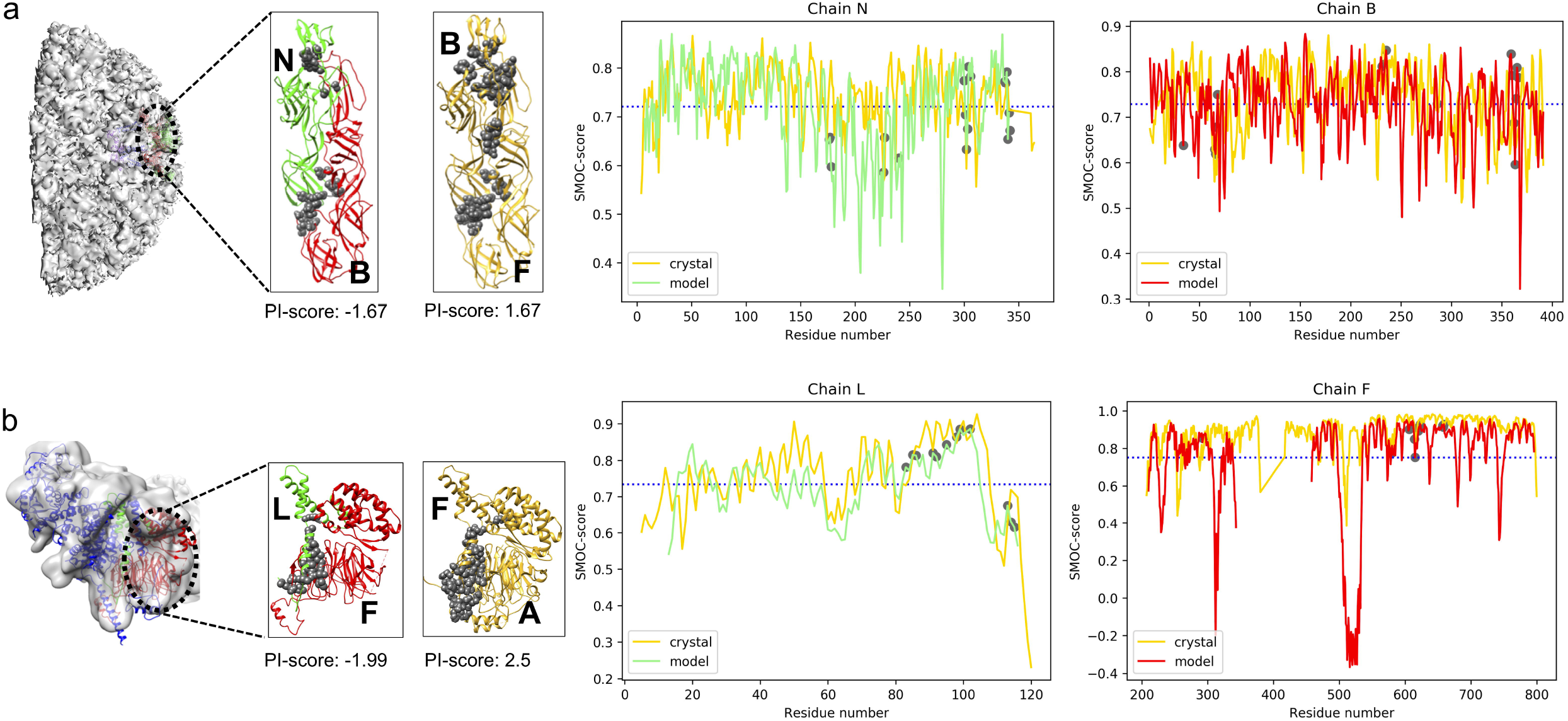
Application to the fitted models in EMDB at intermediate-low resolution. The chains from the crystal structure are in gold and the chains from modelled structure are in red and green. The interface residues are shown as grey spheres. The plot of the local density-based score (SMOC) is shown for the chains forming an interface in the model and the equivalent chain in the crystal structure. The X-axis is numbered as per the residue numbers in the crystal structure. The average SMOC over the model chain is shown as blue dashed line. a) 5 Å resolution structure of Chikungunya virus and the subcomplex envelope1-envelope2 heterodimer (E1-E2) (EMD-5577; fitted PDB: 3J2W). The corresponding 2.35 Å resolution crystal structure is PDB: 3N44 (gold). b) 9.8 Å resolution structure of the TFIID subunit 5 and 9 sub-complex (EMD-9302, fitted PDB: 6MZD, cyan and green). The 2.5 Å corresponding crystal structure for subunit5-subunit 9 is PDB: 6F3T (gold).

We also calculated the density-based scores (global and local) for the E1-E2 subcomplex to assess the fitted model and crystal structure. The E1-E2 subcomplexes from both the fitted model and crystal structure have a CCC of 0.64 and average SMOC over interface residues of 0.72 and hence are indistinguishable with these scores. The plot for the SMOC score (per residue) is shown in Figure 5a, for both chains and the average SMOC score per chain is shown with a blue dashed line. Interestingly, the interface residues (grey circles) are observed to score higher than the per-chain average, especially for chain B. Therefore, at intermediate resolution interface-based scores such as PI-score can prove useful to distinguish the offsets in the modelled protein-protein interfaces that are indistinguishable with the density-based scores.

PI-scores for the interfaces in the fitted models at the intermediate resolution range derived from EMDB are available as Supplementary Table S8.

### Low resolution set (8-12Å)

Transcription initiation factor TFIID complex: A cryo-EM map resolved at 9.8 Å resolution (EMD-9302) with fitted model is available for the structure of the TFIID complex (PDB: 6MZD, shown in green and red in Figure 5b, with interface residues shown as grey circles). The interface between subunits 9 and 5 in the fitted model (LF) was scored negative (PI-score: −1.99). This interface is shifted when compared to the corresponding crystal structure (Supplementary Table S7) at 2.5 Å (PDB: 6F3T, chains F and A, shown in golden with interface residues marked as grey circles, Figure 5b).

Next, we calculated the density-based scores (global and local) to assess the fitted model and crystal structure. The CCC of the fitted model is 0.54 and CCC of the crystal structure is 0.63 upon local optimization of the fit in map, whereas the average SMOC score over interface residues is 0.84 and 0.87 for the fitted model and crystal structure, respectively (Figure 5b). In this example, we see again (but this time with low resolution maps) that PI-score can provide additional complementary assessment when density-based scores alone are not sufficient to identify the offsets in the modelled interfaces.

PI-scores for the interfaces in the fitted models at the low-resolution range derived from EMDB are available as Supplementary Table S9.

### Application to SARS-CoV-2 Cryo-EM Derived Complexes

We also assessed the fitted models in the SARS-CoV-2 cryo-EM maps using PI-score. 108 fitted models were downloaded from EMDB [https://www.ebi.ac.uk/pdbe/emdb/searchResults.html/?EMDBSearch&q=text:(ncov%20OR%20SARS-CoV-2)]. Out of the 108 models, we were able to successfully compute interface features and PI-scores for 55 complexes (149 interfaces). Of these 149 interfaces, 12 were observed to have a negative PI-scores (Supplementary Table S10), with 11 of these being antibody-antibody or protein-antibody interfaces. As our machine learning classifier is not trained on such interfaces (which are reported to have different shape complementarity from other protein-protein interfaces^17^), we decided not to further investigate these cases.

However, the interface between small subunits (S28-S5) of a human 40S ribosome bound to *nsp1* (blue spheres, Supplementary Figure S2a) SARS-Cov2 protein (EMD-11301, PDB ID: 6ZMT) obtained a negative PI-score of −0.04 (Supplementary Figure S2b). We next inspected this complex using the validation suite in CCP-EM (Joseph A.P, *et al*., under preparation). The sub-complex S28-S5 was found to have a clashscore of 7.20 with severe clashes reported at the interface. We used *‘real space refine zone’* and *‘auto fit rotamer’*, with backrub rotamers switched on to fix the steric clashes at the interface using Coot^40^. Upon re-refinement in Coot, the clashscore drops to 6.20 and PI-score improves to 0.25 (Supplementary Figure S2c). The improvement in the PI-scores is most likely due to resolving the clashes between the interface residue pairs R63-A138, V55-34S and L59-R122 (Supplementary Figure S2b) from chain d and K, respectively.

### Comparison with Protein-protein Interface Statistical Potentials

Next, we compared PI-score to the existing protein-protein interface-based statistical potentials (PIE^41^ and PISA^42^) commonly used for protein-protein docking. PIE and PISA scores provide residue and atomic potentials respectively, and we also used a combination (0.1*PISA + (−0.8)*PIE + PISA*PIE) of these, which is reported to perform better in identifying “native-like” complexes^42^. We used the 30% randomly selected test dataset from the entire set (PD1+PD2+ND) to calculate the statistical potentials (combined PIE-PISA score) for the interfaces. Different weights for SVM-based score and statistical potentials were tried ranging from 0 to 1, with an increment of 0.1. For this dataset, the combined statistical potential score alone was unable to discriminate between the complexes from ND and PD2 datasets which are both derived using docking (whereas the PI-score separates these complexes better (Supplementary Figure S3).

## Discussion

### Density independent PI-score to assess modelled assemblies in cryo-EM maps

Machine learning-based methods trained using interface features have proven to be discriminatory in identifying the ‘native-like’ complexes, and are routinely used for protein interface sites and hotspots prediction using sequence and structure-based features^30^. So far, such methods have not been applied to the atomic models derived from cryo-EM data, where errors at the interface are likely. Here, we have developed a density independent metric to assess the quality of protein-protein interfaces in cryoEM derived models (PI-score), using a machine learning-based method trained on interface features. We carefully collated high-resolution crystal structures of the protein-protein complexes and annotated them with interface features, which were further used to train a machine learning-based classifier. In total, 12 features were calculated for 9727 interfaces in our dataset.

In total, 12 features were calculated for 9727 interfaces in our dataset. A 9727*12 vector was used as an input to train a classifier using Random Forest, Support Vector Machines and Neural Networks. Shape complementarity at the interface, which is a well-known feature to discriminate ‘native-like’ complexes^17^, was observed to be the most discriminatory feature (see section “training the classifier”). Using both PD1 and PD2 as positive labels and ND as negative class labels, we were able to achieve a validation accuracy of 86%using a ten-fold cross-validation.

We show that our PI-score can help in identifying native-like fits from a pool of candidate models (see sections “Application to CASP targets” and “Application to EM model challenge targets”).

### Importance of interface validation in cryo-EM maps

Most of the structures (∼95%) derived using cryo-EM have at least two protein chains. Hence, it becomes crucial to model protein-protein interfaces in such structures. With the recent advances in technology, single-particle reconstructions are getting to near-atomic resolution, where the modelling of protein-protein interface is becoming more accurate. However, several complexes in the EMDB are in the intermediate-to-low resolution range. The average resolution achieved in 2019 is still less than 5 Å, where the models are likely to be less reliable, especially at regions with less-resolved density.

Additionally, there are plenty of maps where the nominal reported resolution is high, but the local resolution varies significantly. In EMDB, we have identified 107 interfaces in 54 complexes (at resolution better than 4 Å), 508 interfaces in 171 complexes (at 4-8 Å), and 51 interfaces in 23 complexes (less than 8 Å), with a negative PI-score, implying potential modelling issues at the protein-protein interface. Investigating these cases revealed that the errors at the interface could be of different types, including steric clashes, loose interface packing, smaller interface size and lower shape complementarity (see section on “Application to fitted models from EMDB”).

### Comparison with other scores

As we have demonstrated, PI-score is density-independent and is especially useful to distinguish the native-like interfaces at low-to-intermediate resolution, where density-based scores alone become less informative (see section on “Application to fitted models from EMDB”). Most studies calculate the global CCC, which will not reflect minor changes at local regions of the structure (such as interface regions). Local scores can be more informative in this respect; however, they require a well-resolved density around the interface. PI-score captures different type of information, specifically assessing interface quality, and therefore will have an added value. Therefore, we believe that the combination will be extremely beneficial in guiding model fitting and validation (see section “Application to fitted entries in EMDB”).

### Potential usage of PI-score

PI-score has two key uses: to validate and to aid the modelling of interfaces in cryo-EM derived assemblies. We demonstrated its use as a validation score on the CASP and EM model challenge targets. In addition to these, PI-score can also be implemented as part of the model building/refinement process in software packages, such as CCP-EM^43^ and Scipion^44^, to guide the process of model building and model validation (e.g. as part of the CCP-EM validation suite at https://www.ccpem.ac.uk/download.php). Furthermore, PI-score can also be used to filter solutions and identify ‘native-like’ interfaces from protein-protein docking software, such as Z-Dock, and from software that use multi-component assembly fitting approaches, such as PRISM-EM^45^, IMP^46^ and gamma-TEMPy^47.^

### Summary and Future Directions

In this work we have introduced an interface-centric metric, PI-score, and systematically assessed, for the first time, cryo-EM derived assemblies. We are working towards expanding the features’ set (*e*.*g*. coevolution scores) to calculate PI-score and implementing deep-learning approaches. We believe that PI-score will be a crucial addition to the set of validation scores currently used in the cryo-EM community as part of structure modelling tools. It is likely that many protein-protein interfaces in future-deposited cryo-EM structures will contain errors, especially in low-to-intermediate resolution structures. Scores such as PI-score, which provide insights into interface modelling, have the potential to be extremely beneficial if included in the EMDB validation report.

## Methods

### Dataset of High-Resolution Complexes (Positive Dataset1-PD1, ‘native’interfaces)

High-resolution crystal structures of complexes were obtained from PDB with following filters:

1. Minimum number of chains =2
2. Experimental method = X-ray
3. Resolution between 0.0-2.5Å
4. R factor (all) between 0 and 0.25
5. R-free between 0 and 0.3
6. Length of each chain >= 30 amino acids

Using these filters, we fetched the non-redundant PDB structures at 40% sequence identity, resulting in a total of 3926 complexes.

The complexes were further processed to remove symmetric interfaces present in the same structure using *iAlign*^33^ to structurally align the protein-protein interfaces between different chains of the same PDB structure. At the recommended cut-off of interface similarity score of 0.7, non-identical or identical monomers forming similar interfaces were filtered out and the final set contained 2315 complexes with 2858 interfaces.

### Dataset of Near-Native Complexes (Positive Dataset2-PD2)

“Near-native” complexes were derived from the native complexes (PD1). The pair of interacting chains from PD1 dataset were subjected to protein-protein docking using ZDOCK^34^ and the poses (interfaces) with f_Nal_ (fraction of aligned native interface residues) >=0.7 and iRMSD (interface root mean square deviation)^33^ <=3Å were selected.

### Dataset of Non-biological Complexes (Negative Dataset-ND)

The pairs of interacting chains from PD1 were subjected to docking using ZDOCK and the docked poses with f_Nal_ <0.3 or iRMSD >4Å were selected.

### Interface Assignment

Interface residues between two chains were defined using the distance-based threshold of Cα-Cα distance of 7 Å. An interface was only included in the datasets, if it contained at least ten residues from each of the interacting chains.

### Calculation of Interface Features

The following interface parameters were computed:

1. <u>Number of interface residues</u> (num_intf_residues) This was calculated using an *in-house* python script to assign the interface as explained above and count the number of residues from each chain of the complex.
2. <u>Conserved residues at the interface</u> (conserved_interface) For each chain in a given protein-protein complex, the homologs were collected using *PSI-BLAST*^48^ (number of iterations = 3, e-value = 10^−5^, query coverage=80%) against Swiss-Prot^49^ database. The homologs were further clustered at 90% sequence identity using *usearch*^50^, and subsequently aligned using MUSCLE (v3.8.31)^51^. The conservation scores were calculated using the generated multiple sequence alignment as input to the maximum likelihood-based method *Rate4Site*^52^, which measures the evolution of amino acids residues and identifies functionally important sites. The intersection of conserved residues and interface residues (as assigned above) were selected as a set of conserved residues at the interface.
3. <u>Charged residues at the interface</u> (charged) The charged (Asp, Glu, Lys, Arg) amino acids were counted at the interface and this was normalised by the total number of interface residues.
4. <u>Polar residues at the interface</u> (polar) The polar (Ser, Thr, Asn, Gln, His and Tyr) amino acids were counted at the interface and this was normalised by the total number of interface residues.
5. <u>Hydrophobic residues at the interface</u> (hydrophobic) Hydrophobic amino acids (Ala, Leu, Ile, Val, Phe, Trp, Cys, Met) were counted at the interface and this was normalised by the total number of interface residues.
6. <u>Number of contact pairs</u> (contact_pairs) The contact pairs were defined as the number of atomic contacts between the interface residues from the interacting chains.
7. <u>Shape complementarity (sc)</u> This geometric shape complementarity of protein-protein interfaces were computed using the program-SC^17^ from the CCP4^53^ software suite. The value of the calculated statistic sc (shape correlation) describes the extent of interactions of the two chains with respect to each other and varies between 0-1. Protein-protein interface with sc = 1 suggests that the two protein subunits mesh precisely, whereas with sc closer to zero implies an interface with uncorrelated topography. The following features were calculated using PISA (*via* CCP4) (Protein interfaces, surfaces and assemblies):^54^
8. <u>Hydrogen Bonds</u> (hb) The number of potential hydrogen bonds at the interface
9. <u>Salt Bridges</u> (sb) The number of potential salt bridges at the interface
10. <u>Interface solvation energy (int_solv_en)</u> The difference in energy between the bound and unbound monomers due to solvation effect.
11. <u>Hydrophobic p-value (pvalue)</u> Probability measure of the specificity of a given interface. The lower the probability is, the more specific the interface is.
12. <u>Interface surface area</u> (int_area) Surface area, which becomes inaccessible to the solvent upon interface formation, measured in Å^2^.

### Importance of Interface Features

Five methods (Ridge, Random forest, Recursive feature elimination, Linear Regression, and Lasso) were used to rank the importance of each of the interface features. We used the *sklearn* Python package with default parameter settings. The mean scores from each of these methods were used to rank the derived features.

### Performance Assessment Metrics

True positive (TP), true negative (TN), false positive (FP) and false negative (FN) were used to assess the performance of the model using the following definitions:

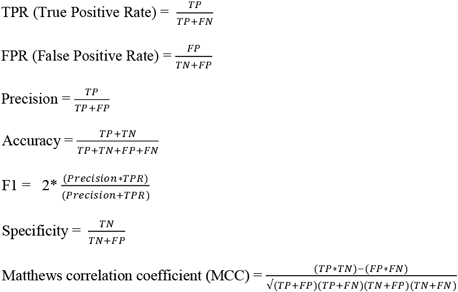

### Benchmark Datasets

1. The structures of fitted models and targets, and corresponding target cryo-EM maps were downloaded from the CASP13 website (https://predictioncenter.org/casp13/).
2. The targets’ structure, map and submitted models for EM model challenge 2016 and 2019 were downloaded from EM model challenge website (https://model-compare.emdataresource.org/).
3. The entries with fitted models were obtained from EMDB (https://www.ebi.ac.uk/pdbe/emdb/).

### Density-based Scores

The goodness-of-fit between the model and cryo-EM map was estimated using global and local cross-correlation scores. The global cross-correlation (CCC) was calculated using Fit-in-Map function in UCSF Chimera^55^ and the local scores (TEMPy SMOC-Segment-based Manders’ Overlap Coefficient^6^) were calculated using the CCP-EM^43^ GUI interface.

### Comparison with CASP13 Oligomeric Scores

The models for CASP13 cryo-EM targets which were scored using our classifier were compared with the protein assembly scores used in CASP13. The machine learning-based classifier score (PI-score) for multiple interfaces within a model structure were averaged by the number of interfaces and this was compared with the CASP13 scores (F1, Jaccard index, lDDT(oligo) and GDT(o))^39^. Interface contact similarity (F1) and interface patch scores (Jaccard coefficient) range from 0 (worst) to 1(best). GDTo and lDDT(oligo) (local distance difference test) consider the whole oligomeric assembly and range from 0 (different quaternary structure) to 1(similar quaternary structure). The latter are computed after mapping the equivalent chains between the target and the model using QS algorithm^39^. If at least one of these CASP13 multimeric scores was >=0.5 and the model was scored positive using our classifier, it was treated as true positive (TP). True negatives (TN) are the set of model structures which do not have any of the CASP13 scores >=0.5 and are scored negative by the classifier. False positives (FP) are the models which were scored >=0.5 by at least one of the four CASP13 scores and negative using our classifier score whereas false negatives (FN) are the models scored negative using the classifier and have at least one of the CASP13 score >=0.5.

### Code Availability

The software to calculate the PI-score is freely available for academic use through: https://gitlab.com/topf-lab/pi_score

## Acknowledgements

We thank Prof. Adrian Shepherd (Birkbeck) for the useful discussions and Dr. David Houldershaw (Birkbeck) for the computer support. We also thank Dr. Andriy Kryshtafovych (UC Davis) for the help and advice with CASP and EM model challenge targets. We are grateful for funding from the Wellcome Trust (209250/Z/17/Z and 208398/Z/17/Z). This work was also partially supported by Wave 1 of The UKRI Strategic Priorities Fund under the EPSRC Grant EP/T001569/1, particularly the “AI for Science” theme within that grant & The Alan Turing Institute.

## Supplementary Files

Supplementary Figure S1: **Scoring the interfaces in the oligomeric target T0995o (CASP13)**. The chains from the labelled structures are marked appropriately and the PI-score is marked for the interface. a. The target structure is shown in golden rod fitted in the cryo-EM density. The dimer interface is scored positive. b. Model TS008_2o, which is the best scoring model in terms of CCC, is superposed onto the target structure. The model is scored positive for the dimer interface. c and d. Model TS117_1o and TS008_5o, respectively, superposed onto the target structure, which is scored negative.

Supplementary Figure S2: **Interface between the small subunits (S5-S28) in the *nsp1*-40S ribosome bound structure (EMD-11301, 3Å)**. a. *nsp1* (blue) bound human 40S ribosome structure, where small subunits S5 and S28 are shown in green and red respectively. b. Close-up of the interface between S5-S28. Interface residues are shown as grey sticks. Steric clashes were observed at the interface and are highlighted in spheres. c. S5(cyan)-S28(pink) upon refinement in Coot. The residues pairs at the interface which were re-modeled are shown as spheres.

Supplementary Figure S3: **Comparison of machine learning based score vs protein-protein interface statistical potentials**. On the X-axis is the combined machine learning statistical potential score (ML_stat_pot combined). The high-resolution complexes (PD1) are shown in yellow, the native-like complexes (PD2) are in green and the negative dataset complexes are shown in skyblue. w1 and w2 are the weights assigned to the machine learning based score and the statistical potentials respectively. a. Performance of two scores are shown for PD2 and ND (negative dataset, with interfaces far from native derived using docking) b.

Performance of two scores for the dataset including native complexes (positive dataset: PD1+PD2 and negative dataset: ND).

Supplementary Table S1: Oligomeric scores for negatively-scoring models using SVM-based model for two of CASP13 cryo-EM targets: T1020o and T0995o. Different scores which assess the oligomeric quality are listed for the models that scored negative using our interface-based SVM score. Interface contact similarity (F1) and interface patch scores (Jaccard coefficient) range from 0 (worst) to 1(best). GDTo and lDDT(oligo) (local distance difference test) consider the whole oligomeric assembly and range from 0 (different quaternary structure) to 1(similar quaternary structure) and are computed after chain mapping combinations between the target to those of the model using QS algorithm (see Methods).

Supplementary Table S2: Density correlation score (CCC) and local score (SMOC) averaged over interface residues for the interfaces in target structure (T1020o) and model (TS208_1o) at target map resolution of 3.3 Å and low-pass filter resolutions (5, 8, 10 and 12Å). For the model TS208_1o, SMOC was not calculated for interface formed by chain B and C (marked as NA) as the number of the residues at the interface was lower than the cutoff.

Supplementary Table S3: Assessment scores for interfaces in the models submitted for three of the CASP13 cryo-EM targets (T0984o, T1020o and T0995o).

Supplementary Table S4: Assessment scores for interfaces in the models scored for the EM model challenge targets.

Supplementary Table S5: Density correlation score (CCC) and local score (SMOC) averaged over interface residues for the interfaces in the target structure (T0002) and model (TS164_1) at target map resolution of 3.3 Å and low-pass filter resolutions (5, 8 and 10Å).

Supplementary Table S6: Assessment scores for interfaces in the fitted models obtained from EMDB at resolution better than 4Å.

Supplementary Table S7: Interface assessment for fitted models in the EM DataBank. The fitted models and crystal structure IDs with their chains identifier forming the equivalent interfaces are listed along with the interface RMSD (iRMSD), fraction of native interface residues aligned (f_Nal_) and the PI-score using our classifier.

Supplementary Table S8: Assessment scores for interfaces in the fitted models obtained from EMDB at resolution range 4-8 Å.

Supplementary Table S9: Assessment scores for interfaces in the fitted models obtained from EMDB at resolution range 8-12 Å.

Supplementary Table S10: Assessment scores for interfaces in the fitted models for SARS-CoV-2 obtained from EMDB.

