## Supplementary material for "Assessment of protein-protein interfaces in cryo-EM derived assemblies": Tables S1, S2, S5 and S7

Table S1:

| **Target** | **Model** | **GDTo** | **lDDT(oligo)** | **Jaccard coefficient** | **F1** |
| --- | --- | --- | --- | --- | --- |
| **T1020o** | TS114_4o | 0.2638 | 0.553 | 0.16 | 0 |
|  | TS008_4o | 0.2212 | 0.529 | 0.09 | 0 |
|  | TS208_5o | 0.274 | 0.568 | 0.14 | 0 |
|  | TS208_3o | 0.2721 | 0.57 | 0.13 | 0 |
|  | TS208_4o | 0.2742 | 0.577 | 0.12 | 0 |
|  | TS208_1o | 0.2734 | 0.567 | 0.12 | 0 |
|  | TS432_3o | 0.0482 | 0.166 | 0.03 | 0 |
|  | TS047_1o | 0.0338 | 0.176 | 0 | 0 |
|  | TS135_3o | 0.2117 | 0.507 | 0.15 | 0 |
| **T0995o** | TS008_5o | 0.3433 | 0.513 | 0.39 | 0.274 |
|  | TS008_4o | 0.2205 | 0.506 | 0.36 | 0.24 |
|  | TS329_1o | 0.2864 | 0.564 | 0.38 | 0.167 |
|  | TS397_3o | 0.2368 | 0.26 | 0.48 | 0.244 |
|  | TS397_4o | 0.3669 | 0.226 | 0.4 | 0.166 |
|  | TS397_2o | 0.3669 | 0.216 | 0.39 | 0.1 |
|  | TS470_3o | 0.1029 | 0.454 | 0.09 | 0.037 |
|  | TS397_1o | 0.2382 | 0.258 | 0.3 | 0.069 |
|  | TS460_1o | 0.104 | 0.59 | 0.24 | 0.022 |
|  | TS196_3o | 0.1081 | 0.579 | 0.1 | 0 |
|  | TS117_1o | 0.103 | 0.559 | 0.16 | 0 |
|  | TS114_5o | 0.0923 | 0.323 | 0.13 | 0.004 |

Table S2:

| Resolution (Å) | Target (T1020o) | | | TS208_1o | | |
| --- | --- | --- | --- | --- | --- | --- |
|  | Chains  Target | CCC  (Chimera) | SMOC | Chains  Model | CCC  (Chimera) | SMOC |
| 3.3 | BC | 0.77 | 0.69 | AB | 0.40 | NA |
|  | BA | 0.77 | 0.69 | AC | 0.39 | 0.27 |
|  | AC | 0.77 | 0.68 | CB | 0.24 | 0.22 |
| 5 | BC | 0.77 | 0.72 | AB | 0.52 | NA |
|  | BA | 0.77 | 0.73 | AC | 0.50 | 0.39 |
|  | AC | 0.77 | 0.72 | CB | 0.38 | 0.32 |
| 8 | BC | 0.79 | 0.73 | AB | 0.68 | NA |
|  | BA | 0.79 | 0.73 | AC | 0.65 | 0.53 |
|  | AC | 0.79 | 0.72 | CB | 0.58 | 0.53 |
| 10 | BC | 0.85 | 0.78 | AB | 0.79 | NA |
|  | BA | 0.85 | 0.77 | AC | 0.76 | 0.63 |
|  | AC | 0.86 | 0.77 | CB | 0.71 | 0.67 |
| 12 | BC | 0.88 | 0.85 | AB | 0.85 | NA |
|  | BA | 0.88 | 0.85 | AC | 0.83 | 0.74 |
|  | AC | 0.88 | 0.85 | CB | 0.80 | 0.77 |

Table S5:

| Resolution (Å) | Alpha-ring interface | | | | Beta-ring interface | | | |
| --- | --- | --- | --- | --- | --- | --- | --- | --- |
|  | CCC  (Chimera) | | Local Score (SMOC) | | CCC  (Chimera) | | Local Score (SMOC) | |
|  | Target  P_Q | Model  F_C | Target  P_Q | Model  F_C | Target  X_Y | Model  n_d | Target  X_Y | Model  n_d |
| 3.3 | 0.85 | 0.73 | 0.78 | 0.23 | 0.85 | 0.65 | 0.79 | 0.26 |
| 5 | 0.89 | 0.83 | 0.84 | 0.39 | 0.89 | 0.73 | 0.83 | 0.51 |
| 8 | 0.93 | 0.91 | 0.93 | 0.57 | 0.93 | 0.83 | 0.93 | 0.71 |
| 10 | 0.94 | 0.92 | 0.95 | 0.67 | 0.95 | 0.89 | 0.96 | 0.80 |

Table S7:

| Fitted model | Crystal structure | Chains of fitted model at interface | Chains of crystal structure at interface | iRMSD (Å), f_Nal_ | PI-score for fitted model |
| --- | --- | --- | --- | --- | --- |
| 3J2W | 3N44 | NB | BF | 2.08, 0.34 | -1.67 |
| 6MZD | 6F3T | LF | FA | 1.68, 0.4 | -1.99 |
