## Supplementary figures and images for "Assessment of protein-protein interfaces in cryo-EM derived assemblies"

### Fig S1

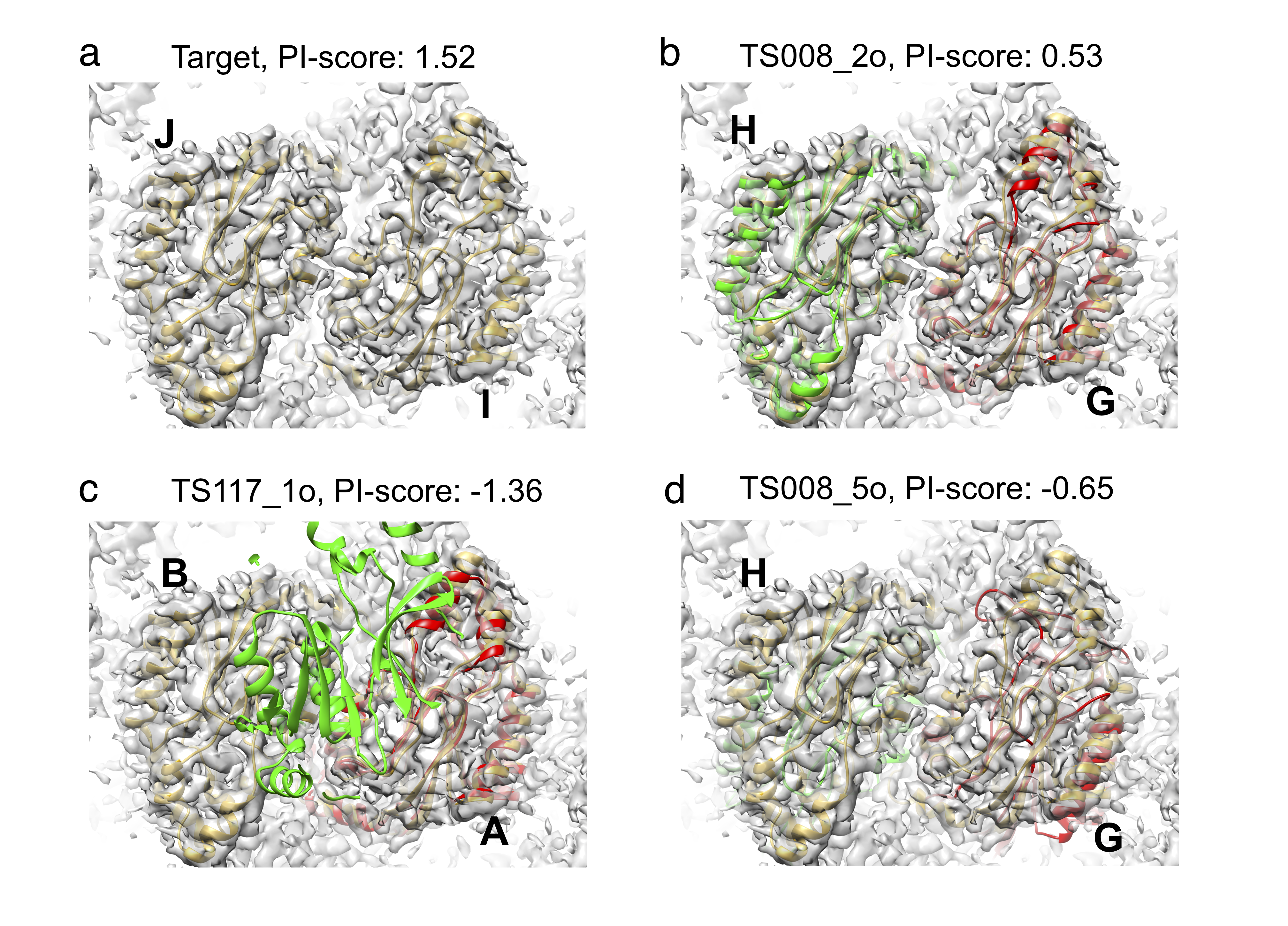

### Fig S2

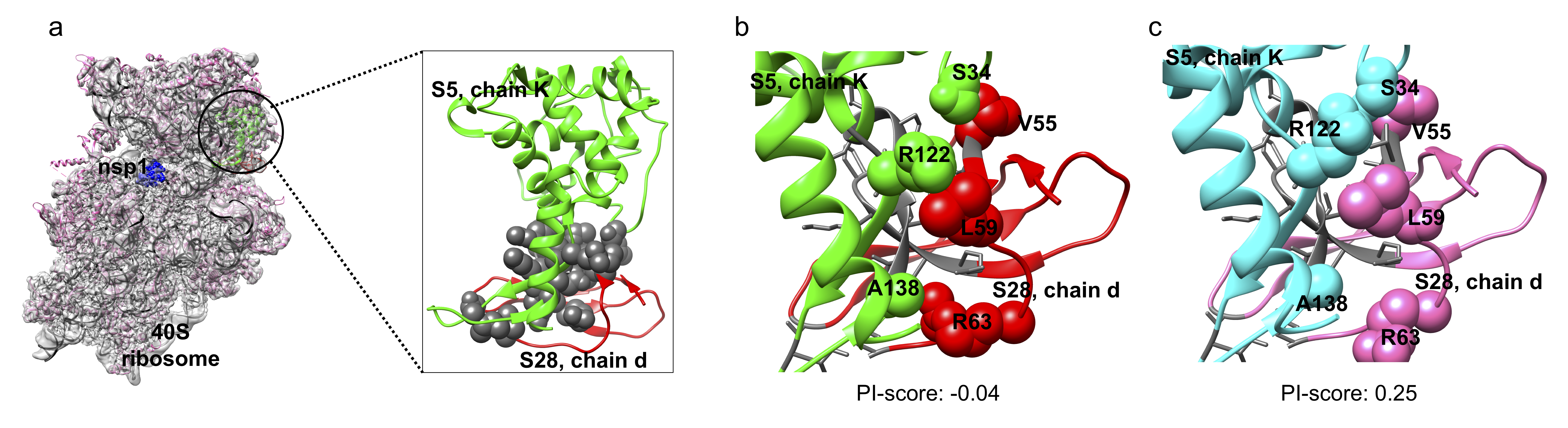

### Fig S3

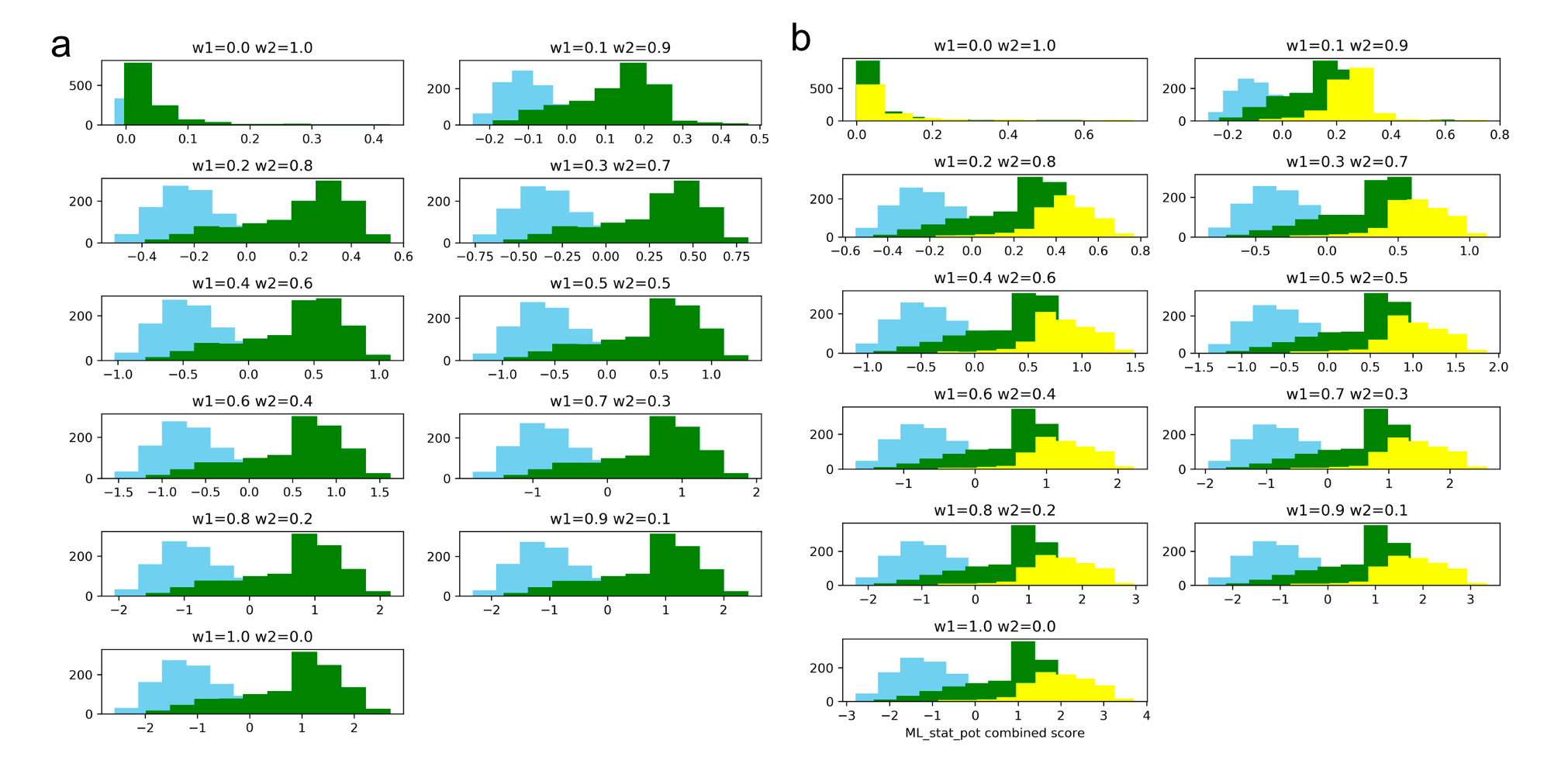
